# NxTrim: optimized trimming of Illumina mate pair reads

**DOI:** 10.1101/007666

**Authors:** Jared O’Connell, Ole Schulz-Trieglaff, Emma Carlson, Matthew M. Hims, Niall A. Gormley, Anthony J. Cox

## Abstract

**Motivation:** Mate pair protocols add to the utility of paired-end sequencing by boosting the genomic distance spanned by each pair of reads, potentially allowing larger repeats to be bridged and resolved. The Illumina Nextera Mate Pair (NMP) protocol employs a circularisation-based strategy that leaves behind 38bp adapter sequences which must be computationally removed from the data. While “adapter trimming” is a well-studied area of bioinformatics, existing tools do not fully exploit the particular properties of NMP data and discard more data than is necessary.

**Results:** We present NxTrim, a tool that strives to discard as little sequence as possible from NMP reads. The sequence either side of the adapter site is triaged into “virtual libraries” of mate pairs, paired-end reads and single-ended reads. When combined, these data boost coverage and can substantially improve the de novo assembly of bacterial genomes.

**Availability:** The source code is available at https://github.com/sequencing/NxTrim

## 1 INTRODUCTION

A common design for DNA sequencing experiments is to sequence from both the 5′ and 3′ ends of the templates in a library to obtain *paired-end reads* for which the genomic distance between the two halves of each pair is approximately known, information which is useful for de novo assembly, alignment and variant calling.

*Mate pair* libraries add further value by increasing the effective genomic distance (EGD) between the two reads. The Nextera Mate Pair protocol is typical: a library of longer DNA molecules is circularised and then fragmented to a size suitable for sequencing, the ends of each circle being joined by an *adapter* sequence tag that is biotinylated to allow enrichment for only those templates that span the join. Read pairs from these templates have an EGD that is determined by the length of the molecule that was circularised, thus yielding longer-range scaffolding information than can be deduced from a standard paired-end read library.

Before further processing, the known adapter sequence must be removed in silico to leave only genomic sequence behind. *Adapter trimming* is a generic task in bioinformatics for which a plethora of tools exist (comprehensively surveyed by Jiang *et al*. (2014)), including some which are specialised to the particular needs of mate pairs (Leggett *et al*., 2013; Jiang *et al*., 2014). However, to our knowledge, all of them work by trimming the adapter and everything to the 3′ side of it, retaining only the portion of the read that lies to the 5′ side of the adapter.

Here we present a tool NxTrim which demonstrates that the 3′-wards portion of the read constitutes valuable “real estate” that can be retained to improve coverage and de novo assembly quality. More specifically, the sequence to the 3′ side of the adapter, together with the other half of the read pair, can be reinterpreted as a standard paired-end read. Depending on where the adapter lies in the read, we reinterpret the whole read pair as a single read plus either a mate pair or a paired-end read, choosing between the latter two options so as to maximise the number of bases that are paired. NxTrim converts raw NMP reads into four “virtual libraries:”

- MP: a set of *known mate pairs* having an outward-facing relative orientation and an EGD whose distribution mirrors the size distribution of the circularised DNA.
- UNKNOWN: A set of read pairs for which the adapter could not be found within either read. Most likely the adapter will lie in the unsequenced portion of the template, although we note (Supplementary Figure 3) some contamination with paired-end reads.
- PE: a set of *paired-end reads*, having an inward-facing relative orientation and an EGD whose distribution mirrors the size distribution of the sequenced templates.
- SE: A set of *single reads*.

Trimming tools following a “5′-only” strategy would produce output similar to the MP and UNKNOWN libraries combined. However, the versatile Velvet de novo assembler (Zerbino and Birney, 2008) can accept all four of these libraries as input to a single assembly and is able to treat the MP and UNKNOWN libraries differently in anticipation of a proportion of non-mate paired reads in the latter.

## 2 METHODS

NxTrim’s logic is described in the Supplementary Materials. Briefly, if the adapter is not found, we place the pair in the UNKNOWN library. If the adapter is detected at the end of one (or both) of the reads, the adapter is removed and the pair is placed in the MP library. If the adapter is at the beginning of a read, the adapter is removed and the pair is placed in the PE library. An adapter in the middle of the read gives rise to a split read. The longest of the split segments is paired with the other read, the pair being added to either the MP or PE library according to which of the 5′-wards or 3′-wards segments is longest. The remaining segment goes into the SE library if its length exceeds a configurable threshold that defaults to 21bp.

We analysed two replicates of each of nine common bacterial samples, all prepared according to the NMP protocol then sequenced as paired 150bp reads during a single run of a MiSeq instrument (Supplementary Table S1). NxTrim’s output was compared to that produced by the MiSeq instrument’s on-board trimming routine, this being exemplary of the “5′-only” approach to adapter trimming employed by all other tools we are aware of.

For each trimmer/sample combination, the reads were assembled by Velvet (version 1.2.10) for all odd *k*-mer sizes between 21 and 119, from which we chose the assembly that maximised contig N50. Contig N50 was strongly correlated with the number of genes detected (Supplementary Figure 2), so this appears to be a reasonable way of selecting an optimal *k*-mer size. We found scaffold N50 to be less correlated with the number of genes found, since it is possible to combine a large number of very small contigs into a long scaffold that has numerous gaps. Assemblies were evaluated using QUAST (Gurevich *et al*., 2013).

## 3 RESULTS

Supplementary Figure 3 shows the insert size distributions of the three virtual libraries, as estimated from alignments to the relevant reference genomes with BWA-MEM (Li, 2013). The MP (centre) and UNKNOWN (right) libraries display pleasingly similar distributions, having median insert sizes of 3.85kbp and 3.69kbp respectively, with a trace of small-insert contamination visible in the latter. The PE library has a median insert size of 299bp.

Assembly comparisons are summarised in Table 1 (with more detail in Supplementary Tables S2 and S3): on average, NxTrim achieves 40.20× coverage, an 18% improvement on the 33.96× obtained by the standard trimming routine. Reads from the standard trimming routine assemble to an average NG50 and NGA50 of 3.025Mbp and 0.993Mbp respectively, while NxTrim improves these metrics to 3.795Mbp and 1.223Mbp. In many cases, the NxTrim assembly has scaffolded nearly the entire bacterial genome. While the lower NGA50 values suggest a number of misassemblies, most of these are due to mis-estimated gap sizes rather than more serious inversions or translocations (as illustrated by the alignments shown in Supplementary Figure 4). We consider NGA50 to be a rather harsh measure here since Velvet does not claim to correctly estimate the gap sizes.

**Table 1.**
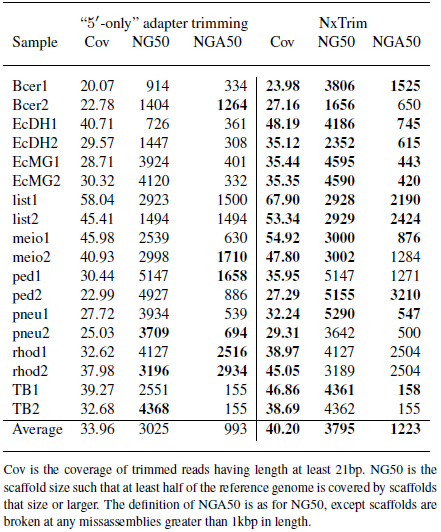
Effect of trimming strategies on coverage and assembly statistics for bacterial samples.

## 4 DISCUSSION

The additional coverage retrieved by NxTrim becomes proportionately larger for longer NMP reads and, while we have used Velvet assemblies to demonstrate its value, should prove equally helpful to other assemblers such as SPAdes (Bankevich *et al*., 2012) and to other applications such as resequencing. Moreover, NxTrim’s ability to extract both long and short insert reads from a single NMP dataset might allow a single mate pair library to suffice for applications where a dual library experimental design might previously have been considered.

### Funding

All authors are employees of Illumina Inc., a public company that develops and markets systems for genetic analysis, and receive shares as part of their compensation.

